# Super-resolution imaging of highly curved membrane structures in giant vesicles encapsulating molecular condensates

**DOI:** 10.1101/2021.08.04.455034

**Authors:** Ziliang Zhao, Debjit Roy, Jan Steinkühler, Tom Robinson, Reinhard Lipowsky, Rumiana Dimova

## Abstract

Molecular crowding is an inherent feature of the cell interior. Synthetic cells as provided by giant unilamellar vesicles (GUVs) encapsulating macromolecules (polyethylene-glycol and dextran) represent an excellent mimetic system to study membrane transformations associated with molecular crowding and protein condensation. Similarly to cells, such GUVs loaded with macromolecules exhibit highly curved structures such as internal nanotubes. In addition, upon liquid-liquid phase separation as inside living cells, the membrane of GUVs encapsulating an aqueous two-phase system deforms to form apparent kinks at the contact line of the interface between the two aqueous phases. These structures, nanotubes and kinks, have dimensions below optical resolution and if resolved, can provide information about material properties such as membrane spontaneous curvature and intrinsic contact angle describing the wettability contrast of the encapsulated phases to the membrane. Previous experimental studies were based on conventional optical microscopy which cannot resolve these membrane and wetting properties. Here, we studied these structures with super-resolution microscopy, namely stimulated emission depletion (STED) microscopy, together with microfluidic manipulation. We demonstrate the cylindrical nature of the nanotubes with unprecedented detail based on the superior resolution of STED and automated data analysis. The spontaneous curvature deduced from the nanotube diameters is in excellent agreement with theoretical predictions. Furthermore, we were able to resolve the membrane “kink” structure as a smoothly curved membrane demonstrating the existence of the intrinsic contact angle. We find very good agreement between the directly measured values and the theoretically predicted ones based on the apparent contact angles on the micrometer scale. During different stages of cellular events, biomembranes undergo a variety of shape transformations such as the formation of buds and nanotubes regulated by membrane necks. We demonstrate that these highly curved membrane structures are amenable to STED imaging and show that such studies provide important insights in the membrane properties and interactions underlying cellular activities.

---

Cells ubiquitously exhibit complex networks of highly curved membrane structures. Examples include the elaborate membrane meshwork of the endoplasmic reticulum, the Golgi body and the inner mitochondrial membrane as well as membrane nanotubes for cellular transport, communication, and motility^1–5^. The dimensions of these highly curved membrane features are typically below optical resolution and pose great challenges for direct live visualization and characterization using conventional microscopy methods. However, emerging super-resolution techniques such as stimulated emission depletion (STED) microscopy^6^ drastically improve the optical resolution limit to the nanometer regime, thus allowing for direct visualization of these highly curved membrane structures. STED microscopy uses two overlapping synchronized laser beams which scan through the sample consecutively, the first laser beam excites the fluorophores in the sample and the second one depletes the excitation everywhere but at the center of the focus, thus restricting the region of fluorescence emission to a size far smaller than the diffraction limited focus of the excitation laser beam^7^. The lateral resolution of STED microscopy generally reaches 20 - 50 nm, which is a drastic improvement compared to conventional confocal microscopy. In addition, one of the biggest advantages of STED over other super-resolution techniques such as photo-activated localization microscopy (PALM) or stochastic optical reconstruction microscopy (STORM) is its unique ability for instantaneous live imaging (at video frequency)^8^. Over the past several years, due to this unique feature, STED has been applied to study biological samples in real time. Examples include imaging of the slow morphing and movements of organelles including endoplasmic reticulum or microtubules^9, 10^, subcellular organization in live cells^8, 11^, and synaptic structures in live samples^8, 12–14^. Thus, it proves to be a promising application for live imaging and measuring membranous structures with sub-optical dimensions. Furthermore, STED imaging of pulled membrane nanotubes was shown to yield direct access to membrane tension^15^.

Different model systems have been used to better understand the mechanics and role of the highly curved membrane structures in cells. Giant unilamellar vesicles^16, 17^ (GUVs) represent a very suitable and minimal cell mimetic model allowing direct microscopy interrogation of the membrane at cell-size scales. Similarly to remodeling cell membranes^18^, membrane tube formation in GUVs can be generated either spontaneously (see e.g. ^19–22^) or via pulling by means of e.g. optical tweezers^23, 24^ or molecular motors^25, 26^. GUV encapsulation of cytoplasmic-like solutions such as coacervates^27^ or aqueous two-phase systems^28, 29^ (ATPS) provide approaches that pave the path towards understanding membrane morphological changes in organelles as well as remodeling by molecularly crowded protein condensates. GUVs encapsulating ATPS of polyethylene glycol and dextran have been already shown to exhibit the formation of nanotubes^30^. Their diameters could be directly measured only for relatively stiff membranes (in the liquid-ordered phase) which form micron-thick tubes, while the radii of thin nanotubes were deduced only from theoretical considerations^31^.

Here, we interrogated this model system with STED microscopy to directly measure the nanotube curvature and thus (i) demonstrate the suitability of STED for quantitative measurements on nanotubes and highly curved membranes, and (ii) probe the validity of the theoretical modeling^31^. The precise imaging of the nanotubes was facilitated by their adsorption to the ATPS interface. Furthermore, phase separation in the GUV bulk and subsequent deflation, have been shown to lead to vesicle budding.^32^ The cross-sectional optical imaging of the bud neck appears to show the presence of a membrane “kink”. Similar “kink”-like structures are observed at membranes of protein storage vacuoles in contact with condensates of seed storage proteins^33^. However, such apparent “kinks” cannot persist on nanometer scale because of bending energy constraints^34^. In this work, STED microscopy allowed us to optically resolve the morphology of the “kink”. This was made feasible by microfluidic trapping of the GUVs, allowing the controlled deflation and imaging of individual vesicles and thus bringing to light the detailed nanostructure of the highly curved membranes with unprecedented resolution. Our experimental findings demonstrate that this unprecedented resolution of STED can easily resolve and quantify membrane shapes in minimal cells (giant vesicles) opening avenues towards resolving the intricate structure of cellular organelles and biomembrane transformations during cellular processes.

## RESULTS AND DISCUSSION

### STED Alignment and Resolution

In STED microscopy, laser alignment is crucial for achieving resolution in the nanometer range. Here, we used 150 nm gold beads for the alignment of the excitation beam with the donut-shaped depletion beam, Fig. 1a. Using TetraSpeck beads of four colors, we also corrected mismatches between scattering and fluorescence modes^35^. In the case of 3D STED, a depletion laser with a phase pattern of a “bottle beam” is enforced in order to improve the resolution in z-axis, Fig. 1b. Compared to confocal imaging, substantial resolution enhancement can be seen in the resulting STED image where more details about the exact location of the beads are observed, Fig. 1c,d.

**Figure 1.**
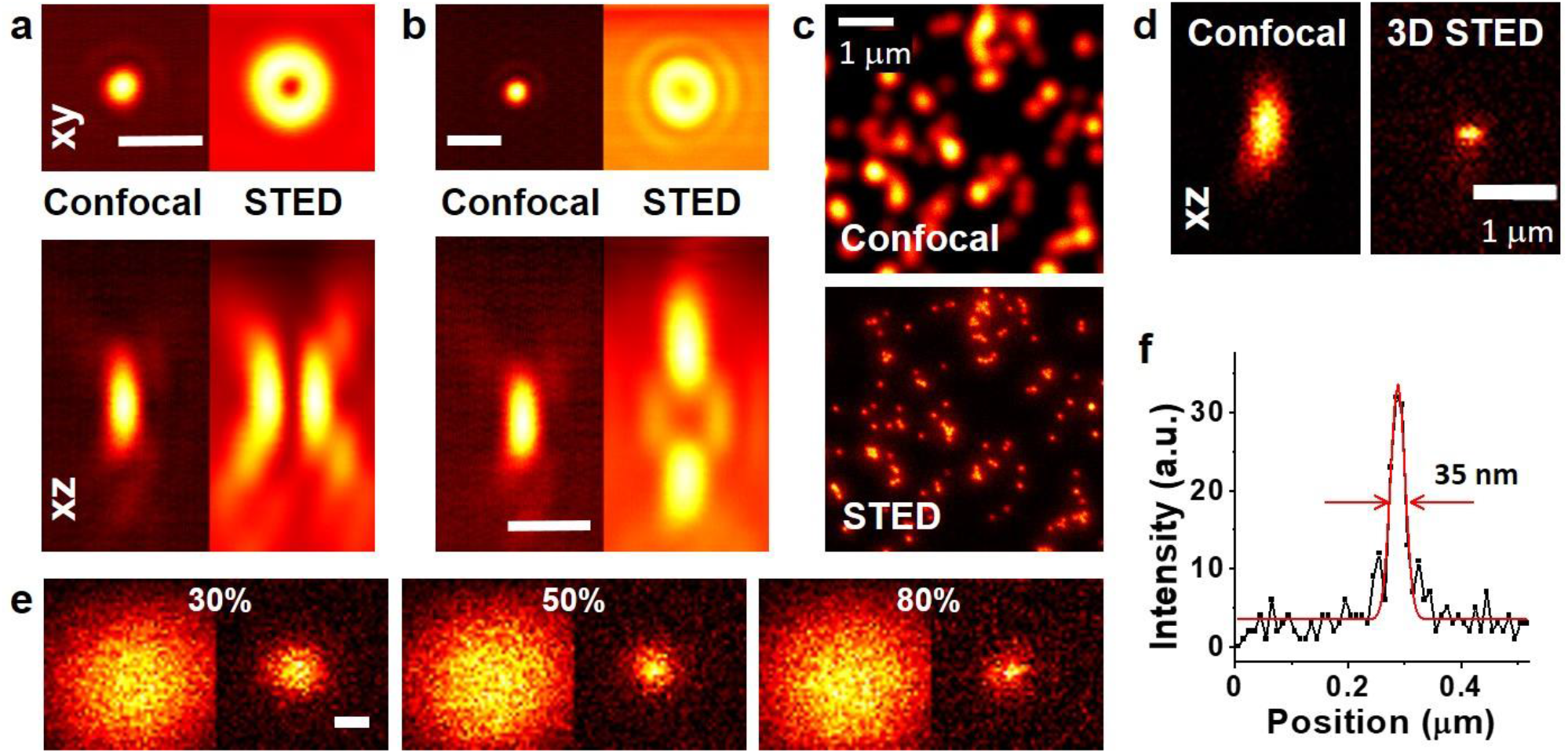
2D and 3D STED alignment and resolution of our system. (a) 2D STED and (b) 3D STED alignment of the excitation and depletion laser beams in xy and xz planes in reflection mode as mapped out by 150 nm gold beads. The excitation beam PSF is aligned to the center of the depletion beam PSF by a spatial light modulation to enhance the resolution. The wavelengths of the excitation and depletion lasers are 640 nm and 775 nm, respectively. Scale bars: 1μm. (c) Confocal and STED images of TetraSpeck beads (100 nm diameter). A clear resolution enhancement can be observed in the STED image if compared with the confocal one. (d) Confocal (left) and 3D STED (right) xz scans of a TetraSpeck bead showing improved axial resolution of around 110-120 nm. (e) Confocal (left) and STED (right) xy scans of a sub-optical crimson bead (26 nm diameter). The bead size appears smaller (approaching its real size) indicating improved resolution with increasing power of the STED laser as indicated on the images in percentage of total power (1.25 W). Scale bar: 100 nm. (f) STED line scan (black curve) of a crimson bead (26 nm diameter) and the respective fitted PSF (red). A FWHM of 35 nm on the bead indicates lateral STED resolution smaller than 40 nm.

Crimson beads of 26 nm diameter which are below the resolution limit of STED microscopy were used to measure the resolving power of our setup. The resolution increases with STED beam power (Fig. 1e). At 80% of the STED laser power (total laser power of 1.25 W), the full width at half maximum (FWHM) of the point spread function (PSF) corresponds to a measured resolution of ~35 nm (Fig. 1f). This represents a nearly 10-fold improvement in the lateral resolution over that of the corresponding confocal laser (at wavelength 640 nm).

### Microfluidic Trapping and Deflation of Individual GUVs

To spatially confine and therefore make the GUVs, membrane tubes and curved structures amenable to STED imaging (which requires immobilization), GUVs were loaded in the traps of a microfluidic device (a modified version of that reported in ^36^). This approach also allows complete solution exchange around the trapped vesicles and thus ensures control on vesicle deflation. The device was initially filled with degassed isotonic polymer solution. Then, a desired amount of GUV stock solution was pipetted on top of the inlet reservoir, letting the GUVs sediment by gravity. A flow (speed of 1.0 μL/min) was applied to load the GUVs into the traps using syringe pump in a suction mode (higher flow rates were observed to deform and damage the vesicles by pressing them against the trapping posts). Side posts in the main channels were used to divert and guide the GUVs pathway into the traps for more efficient filling; see also Fig. S1 in the supporting information (SI) for a sketch of the microfluidic chip. Generally, a large number of GUVs (10-20) could be collected in a single trap of the device, see Fig. S2, but the trapping of individual vesicles was also feasible. Once GUVs were loaded in the device, a gentle flow of 0.035 μL/min was applied to keep them in one spatial location in the trap and facilitate stable imaging. No-flow environment was also tested, but due to Brownian motion and convection GUVs could displace slightly during the imaging process, which was not desired because even slight motion can affect STED imaging at high resolution. To deflate the GUVs, the solution in the chip reservoir was replaced with degassed hypertonic solutions containing increasing amount of sucrose. The flow speed was set to 1 μL/min for 40 min for complete exchange of the external medium outside the GUVs, see Fig. S3 in the SI for the efficiency of solution exchange. After each deflation step, the GUVs were allowed to equilibrate at a flow speed of 0.035 μL/min for 10-20 minutes, which should be sufficient considering the membrane permeability^37^. The microfluidic device offers advantages and represents a big step forward compared to bulk handling of ATPS vesicles (based on simply mixing solutions) as it can trap a large number of GUVs in a single experiment and at the same time exchange the surrounding medium efficiently while following the same vesicle(s). Furthermore, as we will demonstrate further, by modulating the height of the microfluidic device, we were able to influence the orientation of the budded vesicles.

To characterize the vesicle deflation, we introduce the ratio *r* of external osmolarity to initial osmolarity of the solution inside the vesicle. Before deflation (*r* = 1), the GUVs are mainly tense, spherical and without any internal structures. We then deflate the vesicles (*r* = 1.2) by exchanging the external solution around the trapped vesicles with a hypertonic one, see Fig. 2. At this deflation step, the encapsulated aqueous polymer solution is still in the one-phase region, see Fig. S4 for phase diagram of PEG-dextran solutions and for the osmolarity and density conditions of the individual deflation steps. As the GUV shrinks in volume, the excess membrane area is stored in the form of nanotubes protruding in the vesicle interior (Fig. 2, *r* = 1.2). Upon further deflation (*r* = 1.4), the encapsulated polymer solution crosses the binodal into the two-phase region (Fig. S4). Two aqueous phases form inside the GUV, where the PEG-rich phase has a lower density and is located above the dextran-rich phase. The nanotubes formed from the created excess membrane area accumulate at the spherical-cap interface between the two phases to lower the interfacial tension and are visible only under fluorescence microscopy. The nanotubes can diffuse along but are confined to this quasi two-dimensional interface, which allows us to capture them with STED imaging. Further deflation leads to crowding the interface with nanotubes (Fig. 2, *r* = 1.6). To provide best observation of single nanotube, we typically investigate deflation steps that do not lead to overcrowded interface to avoid the proximity of neighboring nanotubes which affect the signal analysis for assessing the tube diameter. A complete single vesicle deflation experiment is shown in Fig. S4d.

**Figure 2.**
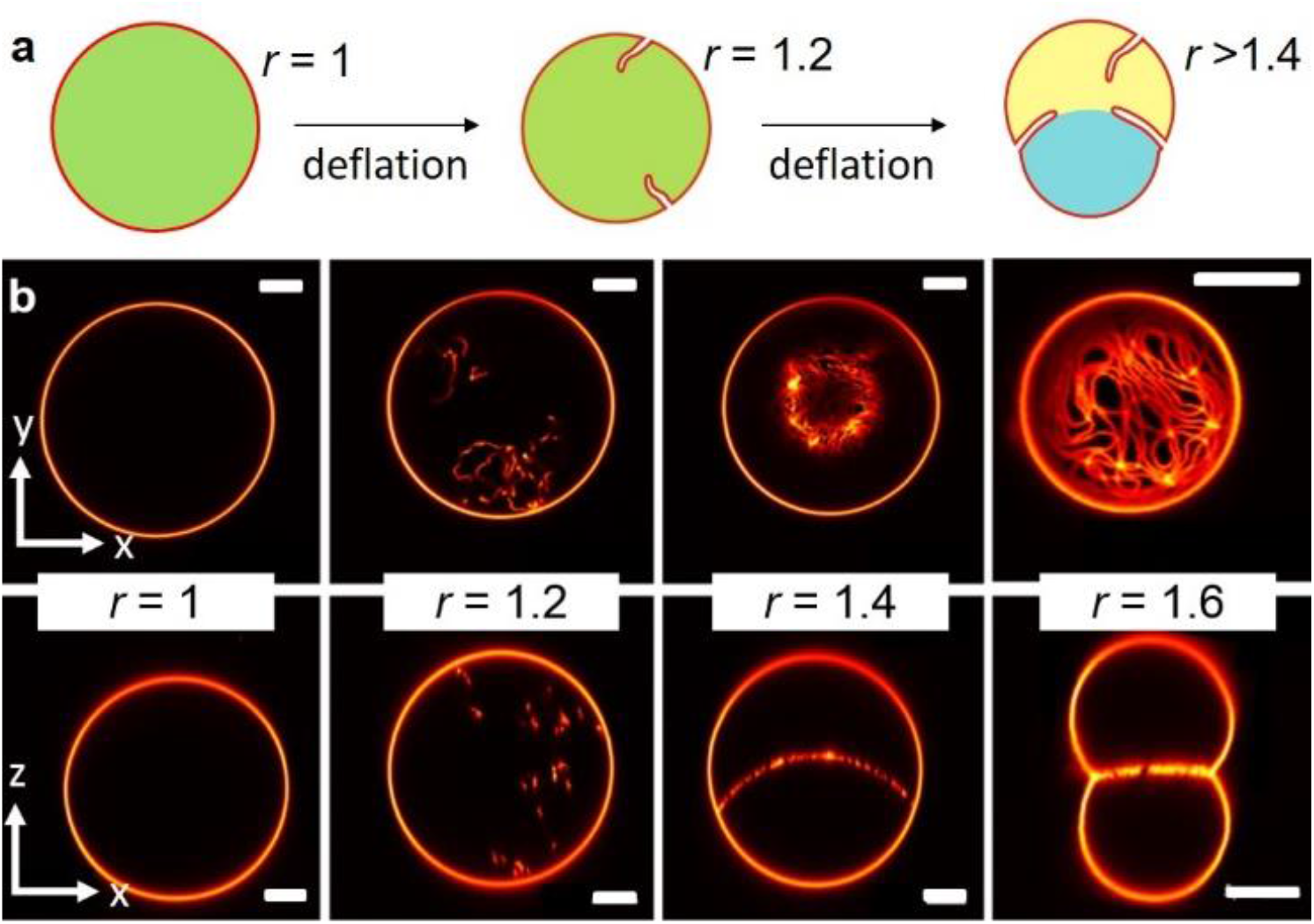
Single-vesicle observation of osmotic deflation, spontaneous tubulation and tube adsorption at the two-phase interface. (a) A sketch illustrating two steps: the GUV initially encapsulates a homogeneous PEG-dextran solution (green) which upon vesicle deflation develops tubes; upon further deflation phase separation into PEG-rich (yellow) and dextran-rich (blue) phase takes place and the tubes adsorb at the interface. (b) Confocal xy scans (upper row) and xz scans (lower row) of morphology transformation of the vesicle for different deflation ratios *r* between the osmolarity of the (introduced) external solution and the initial osmolarity of the solution inside the vesicle; before deflation *r* = 1 (see Fig. S4 for the deflation trajectory in the phase diagram and for additional single vesicle tracking). After the 2nd (*r* = 1.2) and onwards deflation steps, the internal solution in the vesicle undergoes phase separation, the nanotubes adhere at the interface allowing for imaging at a quasi-2D space. The images from the last deflation step (*r* = 1.6) are from another GUV. Scale bars: 10 μm.

### 2D STED Imaging of GUVs and Nanotubes

Confocal images were normally acquired alongside with STED images for the ease of comparison. To acquire a STED image of a full size GUV, normally in the range of ~60 μm, compromises have to be made based on pixel size and image capture time. Choosing a smaller pixel size allows utilizing the full potential of STED resolution but results in very long imaging time in which membrane fluctuation and tube movement will result in artifactual zigzags in the vesicle and tube contours. Therefore, a tradeoff has to be made. Figure 3a,b show images acquired with a relatively large pixel size of 80 nm, i.e. not at the full potential of STED resolution, but still demonstrating the advantage over confocal imaging. Under STED, the GUV membrane appears to be much thinner (Fig. 3a), but because its thickness is only around 4-5 nm, the difference is misleading as it is below the resolution in both cases. However, when comparing the nanotubes (Fig. 3b), we observe a very clear difference between the two imaging modes as individual tubes can be distinguished more clearly. In particular, more details can be resolved when the interface becomes crowded with nanotubes, Fig. 3c,d. Whereas in the confocal image the tubes appear as a smeared homogeneous membrane layer (because the distance between the tubes is below the diffraction limit), under STED we can observe their individual contours.

**Figure 3.**
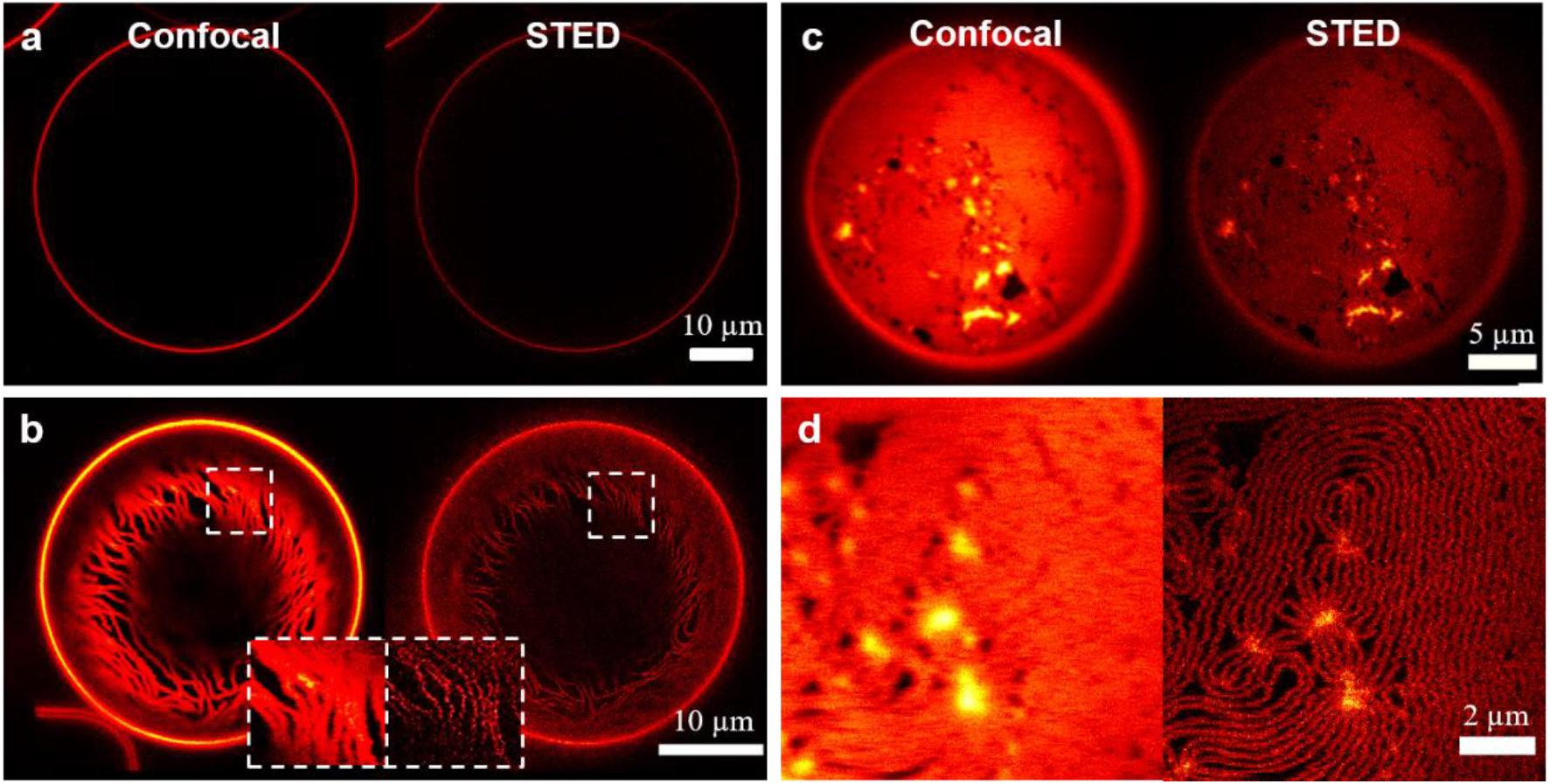
Confocal and 2D STED xy scans of GUVs before and after deflation. (a) A GUV (*r* = 1) trapped in the microfluidic device, no particular difference between the confocal and STED images is observed other than membrane thickness in STED image appears thinner due to brightness difference. (b) A layer of densely packed nanotubes accumulating at the interface of a deflated GUV (*r* = 1.4). Only a fraction of the nanotubes is seen in the scans because the interface is spherical and partially out of focus. Enlarged segments showing the nanotubes are given as insets for better visualization. STED imaging of both (a) and (b) were performed at a pixel size of 80 nm. (c, d) Confocal and STED scans of a densely packed interface with nanotubes after GUV deflation at *r* = 2.0. In contrast to the confocal scans, the individual nanotubes can be clearly distinguished in the STED images. STED imaging of (c) and (d) were performed at a pixel size of 50 nm and 40 nm respectively, to reach a compromise between image acquisition speed and lateral resolution.

After nanotubes adsorb at the interface (and at low deflation ratios), we chose a small scan region (e.g. 4 μm × 2 μm) to image single nanotubes with smaller pixel size (higher resolution). Even though confined to the curved two-phase interface, tubes continue to move due to thermal noise, see Movie S1. As thicker and well-adsorbed tubes move slower, it is feasible to image them with STED and resolve their thickness, see Fig. 4a,b where the left (thicker) tube is less mobile and the right one is partially detached from the interface. While confocal microscopy shows a fuzzy contour limited by conventional lateral optical resolution (in our case ~320 nm), the STED image reveals the tube walls. Intensity profiles from x-direction line scans crossing the tube perpendicularly can be used to measure the tube thickness, Fig. 4c. In this case, the diameter of the left nanotube from the line scans data corrected by nanotube angle of orientation is approximately 150 nm. The thickness appears to be constant along the tube suggesting cylindrical rather than necklace like morphology.

**Figure 4.**
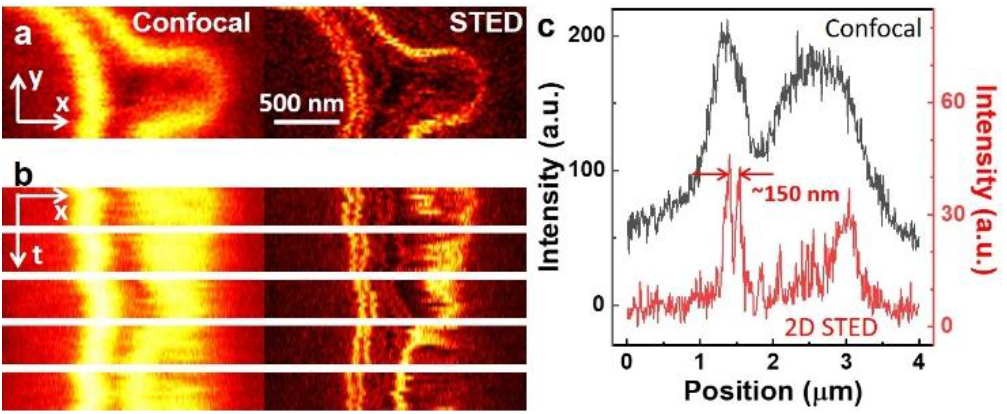
Confocal and 2D STED imaging of two nanotubes at the ATPS interface in a GUV. (a) Confocal and STED xy scans: the left nanotube is well adsorbed and its cylindrical structure is resolved in STED (pixel size 40 nm). (b) Five sets of xt line scans (40 milliseconds in height each); the fuzzy kymograph of the right nanotube shows that it is more mobile (pixel size 10 nm). (c) Example intensity line profiles from confocal and 2D STED line scans in (b). The inter-peak distance in the STED line scan can be used to measure the width of the left nanotube.

### 3D STED Imaging of Nanotubes

Since the nanotubes are fluctuating along the curved ATPS interface, they often get out of focus. Yet, significant signal is collected if fluctuations in z-direction do not exceed the thickness of a confocal slice, typically ~500 nm. According to the Rayleigh criterion, the resolution in z is 1.22*λ*/(*NA*_*obj*_ + *NA*_*cond*_). In our setup, the numerical aperture of the objective is *NA*_*obj*_= 1.2 and that of the condenser *NA*_*cond*_ = 0.55, yielding an axial resolution of 446 nm for our laser at *λ* = 640 nm. Compared to confocal imaging, 2D STED offers no improvement in axial resolution. Thus, we set up the 3D STED alignment for better eliminating the out-of-focus signal in the experiment (see Fig. 1d). Note also that while the diameters of thicker tubes can be resolved with 2D STED (as shown in Fig. 4), for thinner tubes the precision of xt line scans is not sufficient to assess their diameters. Figure 5 illustrates this and provides a comparison of intensity line profiles acquired with 2D and 3D STED on the same nanotube. The two peaks in the 3D STED line profile denote the two wall-crossings of the nanotube which cannot be detected in the 2D STED line scan where light is collected practically from the whole tube cross-section.

**Figure 5.**
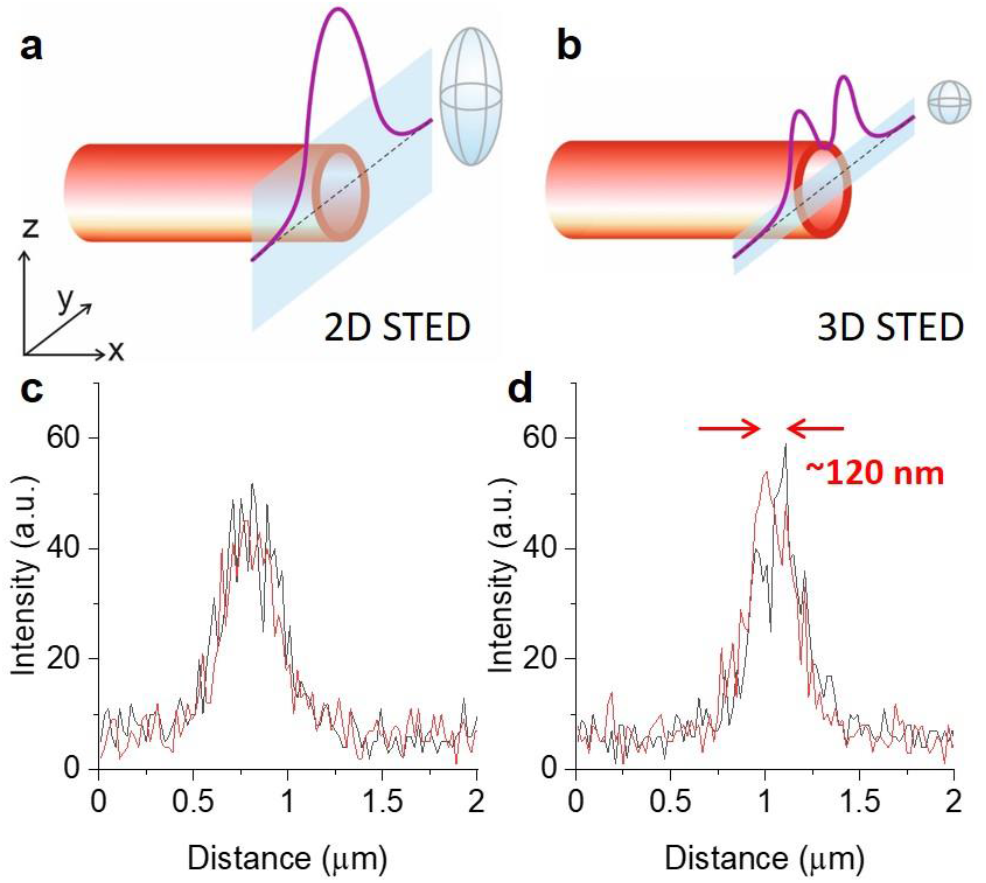
Resolving the diameter of thinner nanotubes with 3D STED. (a, b) Schematic illustrations and (c, d) experimentally acquired data from line scans (gray bands in panels a and b) across a membrane nanotube (red cylinder) when using 2D STED (a, c) and 3D STED (b, d) imaging; for illustration, the approximate dimensions of the scanning voxels are illustrated as gray ellipsoids in (a, b) as well as an idealistic representation of the intensity line scan (purple curves) through the axis of a completely immobilized nanotube. The two intensity line profiles (red and black) in each (c) and (d), were consecutively collected at a pixel size of 20 nm. Contrary to 3D STED line scans, 2D STED line scans do not show two peaks from which the tube diameter can be assessed.

As discussed above, the movement of the tubes hinders the assessment of tube diameters from xy scans. Acquiring single intensity line profiles is however faster. For example, for a pixel size of 20 nm, and pixel dwell time of the laser 10 μs, a complete 4 μm long STED line scan takes 2 milliseconds, while acquiring a 4 μm × 2μm xy scan takes 200 milliseconds, i.e. 100 times slower. In our experiments, we acquired an initial low laser power xy scan to locate the nanotube, followed by a series of (xt) line scans at different locations to deduce the nanotube diameter, and finally another low power xy scan to probe whether the nanotube has displaced (laterally or out of focus) after the line-scan imaging. In the tube diameter analysis, line scans with much lower integral intensity were excluded as they were indicative that the nanotube was no longer in focus.

### Nanotube Shape and Diameter Analysis

The out-offocus tube displacement resulting from thermal noise imposes difficulties in measuring nanotube diameters. Additional uncertainty is introduced if the tube axis is not ideally perpendicular to the line scan direction due to positional fluctuations. All this results in certain error of assessing the tube radius as evidenced from analysis of 3D STED line scans of thicker tubes showing two peaks, see Fig. S5. Most of the manually evaluated diameters (assessed from distance between the two major peaks) fall between 110 and 160 nm for all the deflation levels not showing any trend. On average, around 40 tubes per deflation step were analyzed (see details in Fig. S5). However, 98.2% of the entire collected data were discarded because even at the higher axial resolution provided by 3D STED, the line scans rarely show two major peaks of similar intensity (Fig. S5b), the distance between which could be used to estimate the tube thickness. Note that because of the high resolution, fluctuations down to almost molecular scales are more dominantly represented than in conventional microscopy. Movements of the nanotubes out of the focal plane pose a particular problem, as, if unfiltered, they would lead to an apparently smaller tube diameter. Also fluctuations of the nanotube wall during the line-scan might lead to an over- or underestimation of the nanotube diameter. To improve statistics, a large number of scans (around 15200) were collected imposing laborious effort for manual evaluation.

To enhance reproducibility, emphasize the statistical significance and reduce time for data-analysis, we developed a computational approach to perform automated image analysis and calculation of the nanotube diameters. This approach relies on the detection and discarding of out-of-focus tubes by peak intensity thresholds that were calibrated by extensive measurements on single nanotubes (details are given in the Methods section and in Fig. S6). Figure 6 shows the obtained diameters on a total of 96 nanotubes; the presented results constitute about 10 % of the total data, which were obtained with three repeats per nanotube. The nanotube diameters (at *r* = 1.4) follow a Gaussian distribution which is wider (S.D. of 35 nm) than the distribution obtained for a single vesicle (S.D. of 23 nm, see Fig. S6c). This is indicative of an underlying distribution of the physical properties that define the nanotube radius between individual GUVs; this distribution is convoluted with the statistic uncertainty of the measurement (caused by movement and fluctuations of the nanotubes as further discussed in the Methods section). Variance in GUV-to-GUV nanotube diameters can be caused by small differences in polymer encapsulation efficiency during fabrication (Fig. S3) and maybe even in membrane composition differences. However, together with the high accuracy of the measurement method and large sample size, this uncertainty does not preclude a high statistical significance of the average nanotube diameter (109 nm, ±5 nm S.E., N = 47 for deflation ratio *r* = 1.4).

**Figure 6.**
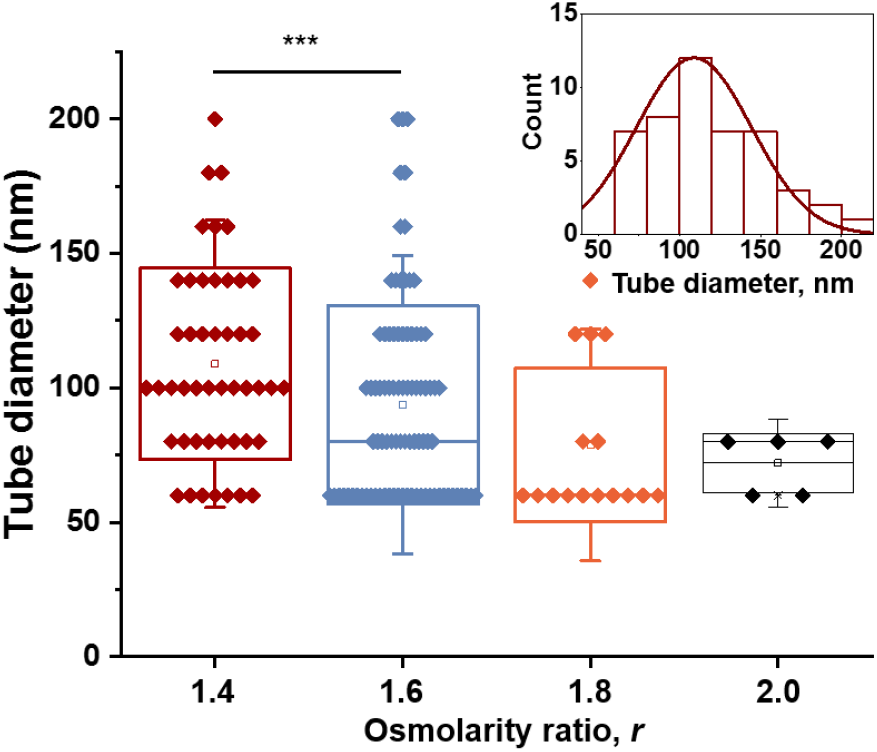
Nanotube diameters measured at different deflation steps using automated analysis. *** indicates p < 0.05 significance estimated from Student’s t-test. The inset shows the histogram of the nanotube diameters measured at the second deflation step (*r* = 1.4) and fit to Gaussian distribution.

### Nanotube Diameter at Different Deflation Levels

One of the aims of this study was, using super-resolution microscopy, to directly probe the validity of previous indirect measurements and theoretical predictions for the membrane spontaneous curvature, *m*, which defines the nanotube diameter, *d*. Previous results suggested a weak dependence of *m* on the deflation levels^31^. We performed a series of direct diameter measurements along the deflation trajectory in our system, reaching deep into the two-phase region (Fig. S4a). With further deflation, we observe a decrease in nanotube diameter, Fig. 6. The diameter distribution at higher deflation levels (*r* > 1.6) are not ideally Gaussian, which is a consequence of the lower tail of the distribution approaching the STED resolution of about 30-40 nm. In addition, at higher deflation ratios, the two-phase interface becomes crowded with tubes (see Fig. 3d) and deflation beyond *r* = 2.0 was not attempted because of overlapping signal (due to tube proximity and movement as observed in the scanning region of 4 μm × 2 μm). Over the whole deflation range, the nanotube diameter varied between 110 and 60 nm.

Note that for the same membrane spontaneous curvature, the diameter of a cylindrical tube *d* = 1/|*m*| is twice smaller than the diameter of a bud *d_b_* in a necklace-like tube, *d_b_* = 2/|*m*|. To deduce whether the tubes had cylindrical or necklace-like morphology, we approximately assessed their persistence length, *L* to be 5.9 μm (S.D. of 3.9 μm) as roughly estimated from the positional dependence of the angle between two tube tangents^38^, see also Movie S2 recorded at *r* = 1.6. For a cylindrical tube of diameter *d*, the persistent length is *ξ_p_* = πκ*d*/(*k_B_T*)^39, 40^, where the membrane bending rigidity is κ ≈ 20 *k_B_T^41^* yielding for *L* between 3.8 and 6.9 μm for tubes with diameters *d* between 60 and 110 nm. On the other hand, the persistence length of necklace-like tubes is comparable to the diameter of the small spheres of the necklace, 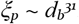. The measured persistence length is close to the one predicted for cylindrical tubes, from which we conclude that the nanotubes are cylindrical. This is consistent with images of very immobile thick tubes as the one shown in in Fig. 4.

### No Influence of Deflation Rate on Nanotube Diameter

It has been previously shown that tube nucleation and growth in stiffer membranes (in the liquid-ordered phase) can lead to coexistence of necklace-like and cylindrical tubes^31^. The transition between these morphologies occurs at a certain critical tube length and is governed by two kinetic pathways that are related to two different bifurcations of the vesicle shape. Upon initial deflation, spherical vesicles can reach the oblate-stomatocyte bifurcation^42^ and exhibit the formation of an inward spherical bud. Further deflation is associated with lipid flow along the bud neck. If this flow is fast, the bud transforms into a growing necklace via sphere-to-prolate bifurcation. If the flow is slow, the vesicle would rather store the excess area by generating more buds. The transition from necklace to cylindrical tube is governed by different contributions (bending energy and volume constraints) and occurs at certain tube length above which the tube has cylindrical morphology^21, 31^. Because of the kinetic origin of the tube shape, we examined whether the rate of deflation in our system would affect the tube diameter measured with STED and whether we can distinguish necklace-like and cylindrical tube morphologies.

The deflation rate was reduced by lowering the flow speed to 0.2 μL/min (5 times slower than in all of the above experiments), while keeping the exchanged volume of medium unchanged and reaching *r* = 1.4. This flow implied that each step of fluid exchange around the vesicles took 200 min (fivefold slower than in the above experiments). The diameter of tubes absorbed at the interface showed no difference compared to those measured at the faster flow rate, see Fig. S7. These results suggest that for the explored flow rates and deflation ratios (1.4 ≤ *r* ≤ 2.0), the shape of the nanotube in our ATPS system is not affected by the deflation speed. Presumably, the tubes were already too long and have undergone necklace-to-cylindrical transformation before adsorbing at the two-phase interface. Indeed, the typical length of the tubes at the deflation ratio *r* = 1.4 was larger than 80 μm (the tubes appear folded at the two-phase interface and their real length can be much longer as only a part of the nanotubes can be imaged on the curved interface). The critical length, above which cylindrical morphology is energetically more favorable, is *L*/*R* > 3, i.e. about three times the vesicle radius *R* ^31^, which is consistent with the observed *L*/*R* values ranging from 4 to 17 for all deflation ratios.

### Membrane “Kink” and Intrinsic Contact Angle Measurement

At deflation ratios above which both the dextran- and the PEG-rich phase partially wet the membrane, the vesicles can adopt budded or snowman-like shapes (see last image in Fig. 2b). Cross sections along the axis of symmetry, misleadingly suggest that the membrane exhibits a “kink” at the three-phase contact line between the PEG-rich phase, the dextran-rich phase and the external deflation medium. Such a sharp “kink”, cannot persist at nanometer range because it will imply infinite bending energy. For this reason, the existence of intrinsic contact angle defining the membrane wetting and geometry was previously postulated^34^. When viewed with sub-optical resolution, the membrane should be smoothly curved rather than exhibiting a kink. Thus, we set to confirm this with direct super-resolution imaging of the bent membrane in this region with STED.

When held in bulk solutions, the GUVs stand upright, because of the density difference in the PEG-rich, external and dextran-rich phases (with increasing density in this order). Confocal xz-scans of such upright vesicles reveals the apparent “kink”, see Fig. S8. Despite the improved 3D STED resolution (~120 nm) in the axial plane (z-direction), it is not feasible to reveal the true structure of the membrane “kink”, Fig. S8. Using a microfluidic device with a controlled height (typically around 60 μm), we were able to orient the budded GUVs to lie with their long axis of symmetry quasi parallel to the coverslip. This allowed us to measure the vesicle geometry from xy scans at the improved lateral resolution of STED. The requirement for precise measurements however, is to image GUVs with two buds which are close in size and comparable to the height of the microfluidic chip, which ensures that the axis of symmetry of the GUV is exactly parallel to the glass coverslip. In this way, we gain direct access to the entire vesicle geometry from a confocal scan through the axis of symmetry. An additional condition is that the two-phase interface is not overly crowded with nanotubes, whose fluorescence signal can obscure imaging of the “kinked membrane”. Taking all this into account, we were able to resolve the precise morphology of the “kink” as a smoothly curved membrane segment, see Fig. 7a-c.

**Figure 7.**
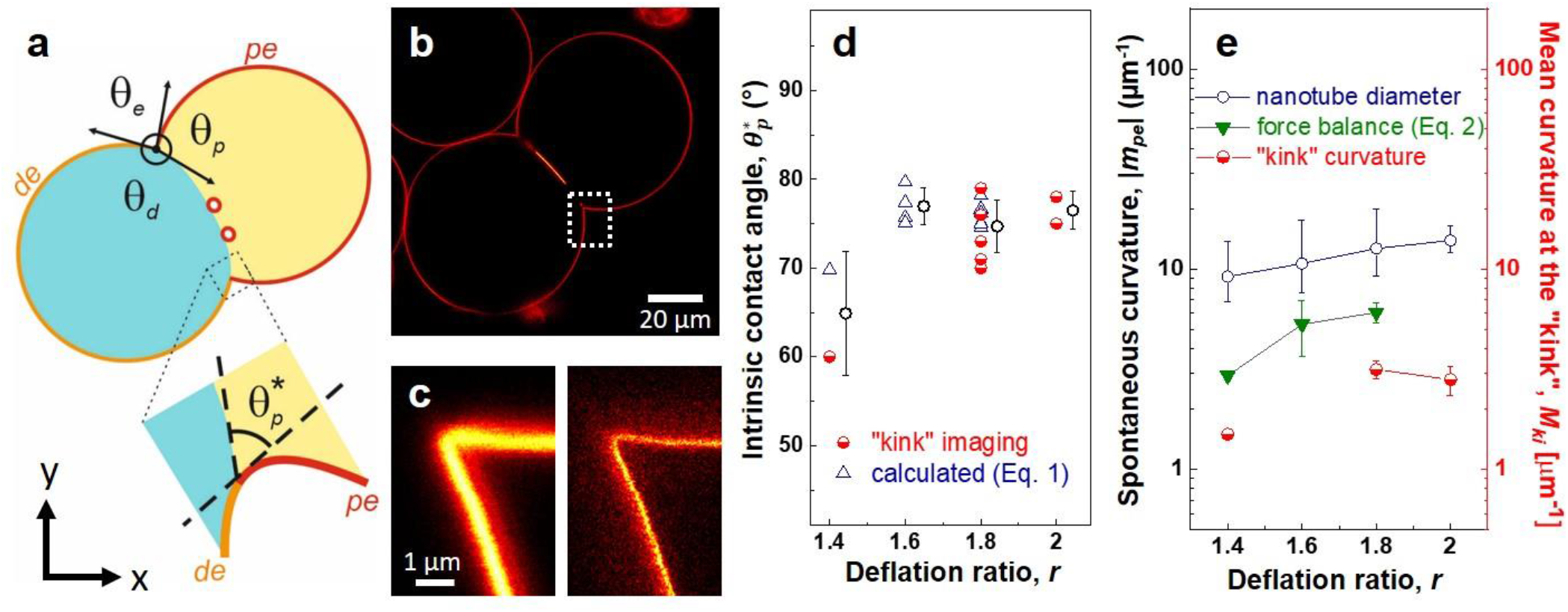
Measuring the intrinsic contact angle and membrane spontaneous curvature. (a) Sketch of a horizontally oriented GUV with a magnified region at the three-phase contact region identifying the apparent contact angles *θ*_*p*_, *θ*_*d*_ and *θ*_*e*_ and the intrinsic contact angle 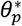. The PEG-rich phase (yellow) wets the *pe* membrane segment (red), while the dextran-rich phase wets the *de* membrane segment (orange). (b) STED image of the whole vesicle (confined laterally in the microfluidic channel) from which the three apparent contact angles are measured, see also Fig. S9. (c) Magnified confocal (left) and 2D STED (right) xy scans in the region outlined by the dashed-line rectangle in (b). The lateral confocal resolution is still insufficient for revealing the morphology of the “kink”, while a smoothly curved membrane with a resolvable curvature is revealed by STED. (d) Values of the intrinsic contact angle at different deflation ratios as estimated from direct STED imaging of the “kink” region at the three-phase contact line (red half-filled circles), and as calculated from Eq. 1 (open blue triangles); mean values from both methods together are shown (black open circles). For deflation ratios of *r* = 1.4 and *r* = 2.0, the sample statistics is poor because (i) membrane undulations render the “kink” radius measurement unreliable, (ii) at *r* =1.4, in most of the cases vesicle budding has not taken place which hinders the readout of the apparent contact angles, and (iii) at *r* =2.0, the two bud centers are no longer in the same plane rendering the measurements of the apparent contact angles and “kink” radius unreliable. (e) Membrane spontaneous curvature, *m_pe_*, deduced from tube diameter |*m_pe_*| = 1/*d* (blue open circles) and force balance Eq. 2 (green triangles), compared to the mean curvature of the membrane at the three-phase contact line, i.e. the “kink” region, *M_ki_* (red half-filled circles).

The geometry of the vesicle in the three-phase contact region is described by the apparent (optically observable) contact angles *θ*_*p*_, *θ*_*d*_ and *θ*_*e*_, see sketch in Fig. 7a for definition. At a fixed deflation ratio, these contact angles may vary from vesicle to vesicle depending on the initial excess area of the individual vesicle in the beginning of the experiment. However, these contact angles can be used to assess the intrinsic contact angle, 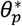, which is a material parameter characterizing the affinity of the PEG-rich aqueous phase to the membrane^34^ (the intrinsic contact angle describing the affinity of the dextran-rich phase to the membrane 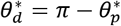:

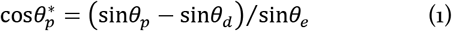

Thus, while the apparent contact angles may vary from vesicle to vesicle at a fixed deflation ratio, the intrinsic contact angle should be the same. We measured *θ*_*p*_, *θ*_*d*_ and *θ*_*e*_ by fitting circles to the vesicle contours at the interface (see Fig. S9a), and calculated the intrinsic angle. The value of the intrinsic contact angle obtained from these apparent contact angles can be compared with the one directly measured from STED images of the area of the contact zone, see Fig. S9b. Using these two approaches, indirect (as theoretically predicted from the apparent contact angles using Eq. 1) and direct (from the high-resolution image of the “kink”), on a series of GUVs at different deflation levels, we see a strong agreement between the theoretical prediction and the direct visualization, Fig. 7d. Our measurements, relying on the dramatic resolution increase by STED, provide the first direct visualization proof of the smoothly curved membrane “kink” and the existence of the intrinsic contact angle. The data also show that at a fixed deflation ratio, the intrinsic contact angle is constant over vesicles with different geometry (Fig. 7d) further emphasizing that it is indeed a material parameter of the system.

We should note that Eq. 1 was derived^34^ under the simplified assumption that the membrane spontaneous curvature is negligible. It is thus somewhat surprising that this simplified expression provides a reliable approximation of the intrinsic contact angle even in the presence of non-zero spontaneous curvature (evaluated in the following section).

Over the whole deflation range (at least at *r* > 1.4), the intrinsic contact angle does not seem to alter significantly attaining an averaged value of ~75°. This value is a little higher compared to previous reports where it was predicted from the apparent contact angles^31, 34^, which is presumably due to the different batch of polymers used.

### Membrane spontaneous curvature

The membrane nanotubes initially form from the membrane segment in contact with the PEG-rich phase and their shape is stabilized by the membrane spontaneous curvature of the *pe* membrane segment, see Fig. 7a. This membrane spontaneous curvature can be assessed from the nanotube diameter, *m_pe_* = 1/*d*, and from the force balance at the three-phase contact line^31^:

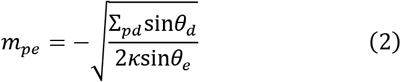

Data for the spontaneous curvature obtained in these two different ways is shown in Fig. 7e. The agreement is relatively good (same order of magnitude) considering the inaccuracy of the bending rigidity value used for the estimates*^41^*. The results obtained from the tube diameter give somewhat higher spontaneous curvature, which could be an indication for (i) a difference between the membrane composition in the tubes and in the *pe* membrane segment of the vesicle body resulting for example in different value of the membrane bending rigidity (for example due to lipid sorting), and (ii) the tubes are adsorbed at the two-phase interface, i.e. in contact with the dextran-rich phase, and thus might be affected by the spontaneous curvature of the *de* membrane segment. From the structure of the “kink” resolved by STED, we can determine the “kink” radius *Rki* in the three phase contact region (see Fig. S9). If we assume that this region is dominated by the curvature of the *pe* segment, i.e. it is a uniform membrane, we can compare the mean curvature of the membrane in the “kink” to the obtained spontaneous curvature. The mean curvature of this toroidal membrane segment is *M*_*ki*_ = (1/*R*_*ki*_ − 1/*R*_*ne*_)/2, where *R_ki_* is the “kink” radius (see Fig. S9) and *R_ne_* is the neck radius of the budded vesicle (i.e. the neck radius of the snowman shape, which contributes only negligibly to the mean curvature). The radius of the “kink” *Rki* ranges from 140 nm to 200 nm (note that due to Brownian noise, the membrane is constantly undulating as shown in Movie S3, which results in substantial scatter of the measured “kink” radius).

The mean curvature in the “kink” region *M_ki_* is compared to the deduced the spontaneous curvature of the membrane in Fig. 7e. The discrepancy could result from the simplified assumption that the curvature in the “kink” region is dominated by the spontaneous curvature of the *pe* membrane segment. It remains to be shown, using even higher-resolution techniques, what the precise morphology and the governing factors shaping the membrane at the nanometer scale are. In any case, the highly curved membrane structures studied here with super resolution microscopy represent stable and transient biomembrane morphologies occurring during various cellular events and exhibited by membrane-bound organelles, and will certainly serve to deepen and expand our understanding of the cell membrane.

## CONCLUSIONS

In this work we use STED in combination with a microfluidic approach to study the highly curved membrane structures in GUVs encapsulating crowded solutions exhibiting phase separation similar to biomolecular condensate formation in cells. The unprecedented super-resolution imaging revealed the cylindrical structure of the liquid-disordered membrane nanotubes for the first time. The spontaneous curvature of the membrane deduced from the diameter of the nanotube is consistent with previous theoretical estimates based on force balance. Furthermore, we prove that the spontaneous curvature of the membrane is only weakly affected by the change of area-to-volume ratio of the vesicle upon deflation.

The movement and fluctuations of the lipid bilayer nanotubes in our experiment posed a certain uncertainty of the measurements due to the finite acquisition time typical for the scanning imaging of STED microscopy. We developed a simple image analysis routine that minimizes this uncertainty and allows for highly accurate nanotube diameter measurements, in principle exceeding the STED resolution by repeated sampling of the same nanotube, and statistical analysis with sub-optical resolution accuracy. Note that compared to other super-resolution techniques (PALM, STORM) the thermal fluctuations and movement pose an even more serious challenge as they require long integration-times on the timescale of minutes, incompatible with the dynamic movements of nanoscopic materials in aqueous solutions. Thus we believe that the approaches developed here will also find applications in other fields of nanoscopy.

The superior STED resolution combined with microfluidic-based positional immobilization and orientation of GUVs allowed us to resolve that the membrane “kink” represents a smoothly curved membrane in the three-phase contact zone corroborating previous theory. Finally, the intrinsic contact angle which only depends on molecular interactions is calculated and compared with the directly visualized cases, showing an excellent agreement with theory (Fig. 7d). The results suggest that the simplified expression (Eq. 1) which ignores spontaneous curvature contributions correctly predicts the intrinsic contact angle for membrane exhibiting spontaneous curvature in the range 5-10 μm^−1^.

The remarkable membrane structures discussed above all share one common character which is a high curvature of the membrane, their generation and evolution upon external stimuli are associated and represent similarities with cellular organelle morphologies that involve shape transformation of the biomembrane. In-depth investigation of these membrane structures based on super-resolution microscopy, as shown here, will pave the way for understanding cellular processes.

## METHODS

### Materials

Dextran from Leuconostoc spp (Mr 450-650 kg/mol, batch number: BCBR8689V), fluorescein isothio-cyanate-dextran (average mol wt 500 kg/mol, batch number: SLCC4853) and poly(ethylene glycol) (PEG 8000, Mv 8 kg/mol, batch number: MKBT7461V) were purchased from Sigma-Aldrich. 1, 2-dioleoyl-sn-glycero-3-phosphatidyl-choline (DOPC) and 1, 2-dipalmitoyl-sn-glycero-3-phosphatidylcholine (DPPC) were purchased from Avanti Polar Lipids and cholesterol from Sigma-Aldrich. 1, 2-Dioleoylsn-glycero-3-phosphoethanolamine labeled with ATTO 647N (ATTO 647N DOPE) was purchased from ATTO-TEC GmbH. All chemicals and lipids were used as received without further purification. All other reagents were of analytical grade. All solutions were prepared using ultrapure water from SG water purification system (Ultrapure Integra UV plus, SG wasseraufbereitung) with a resistivity of 18.2 MΩ.cm.

### Binodal, Critical Point, Density and Osmolarity Measurement

#### Locating the binodal

Cloud point titration was used to determine the binodal and critical point of the mixed dextran and PEG aqueous solution at 23.5 ± 1 °C as previously reported^43^. Concentrated dextran and PEG stock solutions (10-20% by weight fraction) were prepared by dissolving polymers in water. A certain concentration of dextran (PEG) solution was prepared by adding water to the stock solution in a 10 mL sealed glass vial, then PEG (dextran) solution was titrated dropwise into the vial followed by handshaking. The process continued until the thoroughly mixed solution became turbid. The mass of the two stock solutions and water was measured with a balance (Mettler AT 261 Delta Range Analytical Balance) along the process in order to construct the binodal, see Fig. S4a.

#### Locating the critical point

At the critical point, the volumes of the coexisting phases become equal as one approaches the binodal^44^. The critical point of the system was located with a reversed process as described above, namely, approaching the binodal from the two-phase region: A series of solutions in this region were prepared with certain weight ratios between dextran and PEG in a sealed measuring cylinder. Certain amount of water was added into the system followed by vigorous shaking to ensure good mixing, depending on the weight fraction location of the system in the binodal, the system was left to equilibrate for hours or days (more time is required when approaching the critical point) to complete the phase separation process. After the mixed solution finally became clear with an interface in between, the volumes of the coexisting phases were measured.

#### Adjusting the density and osmolarity measurements

In order to balance the osmolarity across the lipid membrane and then further deflate the GUVs, the osmolarity of the mixed polymer solutions without sucrose and with sucrose of increasing concentrations was measured by an osmometer (Gonotec Osmomat 3000 Freezing point osmometer). The densities of these solutions were measured by a density meter (Anton Paar DMA 5000M). A weight ratio of dextran:PEG = 1:1 was chosen as the starting point of the external deflation medium so that the density of the external medium is always in between the densities of the PEG-rich and Dextran-rich phase in the GUV along the deflation trajectory, therefore the GUV was oriented with symmetry axis perpendicular to the coverslip after budding ^21, 45^. This condition facilitates imaging and quantitative microscopy characterization of the system.

### Vesicle Preparation and Deflation

Giant unilamellar vesicles in liquid-disordered phase (Ld) with lipid composition DOPC:DPPC:cholesterol = 64:15:21 were labeled by 0.5 mol % ATTO 647N DOPE. The molar ratio of the dye was chosen so that adequate STED signal can be detected during the experiment. The GUVs were prepared by electroformation as described elsewhere^31^. Briefly, 2-4 μL lipid solution were spread onto each of the conducting sides of two indium tin oxide (ITO) glasses with a Hamilton glass syringe and dried under vacuum for 2 h. The plates were assembled into a chamber with a Teflon spacer and the swelling solution (1.9 mL) of dextran and PEG with initial weight ratio of dextran:PEG = 1.57:1 (4.76%, 3.03% weight fractions) was introduced (this solution gets encapsulated in the GUVs). An alternating electric field of 1.0 Vpp and 10 Hz was then applied using a function generator for 2 h. Afterwards, the GUVs were collected and used immediately. They were dispersed into an isotonic polymer solution with lower density (dextran:PEG =1; 3.54%, 3.54% weight fractions) to facilitate their sedimentation. Vesicle deflation was controlled via a NeMESYS high precision syringe pump (CETONI GmbH) by exchanging the external medium with a series of different hypertonic solutions containing constant polymer weight fractions and an increased weight fraction of sucrose. As indicated above, the density and osmolarity of the external medium were chosen so that the GUVs can not only sediment on the bottom but also “stand” on the coverslip after budding. In certain experiments, where GUVs exhibited budding, to measure the contact angles a microfluidic device with a smaller height (60 μm) was chosen. Each deflation step takes around 40 minutes at a flow speed of 1 μL/min to ensure at least 10 times exchange of the internal volume of the microfluidic device (~4 μL). The successful encapsulation of PEG and dextran aqueous two-phase system in the GUVs was tested by replacing 0.5% weight fraction of dextran molecules with FITC-dextran (molecular weight 500000 g/mol) while keeping the solution conditions the same as normal experiments. The fluorescence intensity inside the GUVs (more than 300 GUVs) was compared with that of the bulk ATPS solution. The solution exchange in the microfluidic device was checked in the vicinity of trapped GUVs. GUVs in the fabrication solution with FITC-dextran both inside and outside were initially trapped, the fluorescent intensity at random positions in the device was acquired. Then, deflation medium without FITC labeled dextran was introduced into the device at a flow speed of 1 μL/min for 40 min and the fluorescent intensity at different places in the device was again checked with the same experimental conditions. The solution exchange efficiency was ~100% as shown in Fig. S3c. During GUV stabilization and imaging, a very tiny flow speed of 0.035 μL/min was chosen so that the GUVs can be slightly pushed onto the posts to prevent vesicle movement yet without obvious morphology change.

### STED Microscopy

#### 2D and 3D STED Alignment and Resolution

STED microscopy uses two overlapping synchronized laser beams which scan through the sample consecutively, the first laser beam excites the fluorophores in the sample and the second one depletes the excitation everywhere but at the center of the focus, thus restricting the region of fluorescence emission to a size far smaller than the diffraction limited focus of the excitation laser beam ^6, 7, 46^. In order to realize this concept for microscopy observation, the point spread function (PSF) of the depletion beam should have zero intensity in the center and maximum intensity at the periphery. This is achieved by placing a liquid crystal spatial light modulator (SLM, usually pixelated liquid crystal devices that change the optical path length by modifying the effective refractive index of each pixel) in the optic path of the depletion laser which forms the desired doughnut-shaped depletion focus pattern. The quality of the doughnut phase pattern from the depletion beam overlapping with the excitation beam determines the ultimate resolution of STED. Scanning the co-aligned beams through the sample yields images of which the lateral resolution is tuned by the intensity of the STED laser^47^.

The STED microscope used here is equipped with a pulsed STED laser beam of 775 nm from Abberior Instruments GmbH. In order to have STED performance at full potential with best resolution, alignment of the excitation and depletion beams is needed. Alignment can be performed by inspection of the depletion focus shape by scanning gold beads near the focus in a reflection mode and adjusting the two focuses until the center of the depletion beam overlaps with the center of the excitation focus. An SLM is used for changing the phase pattern and position of the STED beam. STED laser beam power (total power = 1.25W) was adjusted to 0.04% to protect the detector (pixel size 20 nm, dwell time 10 μs, 1 Airy unit). The parameters in the SLM software for the STED beam were optimized to achieve a good doughnut-shape with zero intensity in the center and uniform maximum intensity along the doughnut ring in xy scan, and the excitation beam focus was positioned to be right in the center of the STED beam so that the fluorophores can be de-excited efficiently. The use of the SLM means that it is straightforward to switch between 2D and 3D STED modes without the relative misalignment that might occur in other systems where phase masks are physically exchanged. 3D STED mode was implemented using SLM to create a so-called “bottle beam” phase pattern in which a zero intensity center is surrounded in all directions by higher intensity^48, 49^. This can simultaneously increase the resolution in the focal plane and along the optical axis. Corrections for mismatches between the scattering mode and the fluorescence mode^35^ was performed using TetraSpeck beads of four colors (TetraSpeck™ Microspheres, 100 nm, fluorescent blue/green/orange/dark red) to check the STED alignment in fluorescence imaging mode. Crimson beads of 26 nm diameter (2% Solids, Carboxylate-Modified Microspheres, FluoSpheres™, Molecular Probe) with size below the STED resolution were used to measure the resolution of STED microscopy. Beads solution were sonicated for 30 min, then diluted 100000 times with ethanol. An aliquot of 2 μL beads solution was deposited on a cover slip and dried under vacuum for 30 min.

#### STED Experimental Conditions

GUVs were observed by a STED microscope (Abberior Instruments GmbH) equipped with 60× Olympus UPlanSApo water immersion objective (N.A. = 1.20). STED images of full size GUVs were captured with STED power of 80% and pixel size between 80-100 nm to have short image acquisition time and stable images. In order to measure the moving nanotubes, confocal beam power was set at 40% and STED at 80% to obtain a substantial STED signal for xt line scans. The xy scanning range was decreased to 4×2 μm, and pixel size to 40 nm with confocal laser power at 10% to avoid bleaching the nanotubes. The xt line scan length was set at 4 μm and pixel size to 10-20 nm according to Nyquist theorem for correct data sampling to have full potential of STED performance based on its resolution. During scanning and data acquisition, an initial xy scan was executed on a single nanotube, then a series of xt line scans were performed on different locations of the nanotube along the y axis. A final xy scan was collected to show the final position of the nanotube and whether it has remained in the scanning range. Each line scan took about 2 milli-seconds and the whole xy and xt series took less than 1 second. Since the nanotubes exhibit constant (mainly lateral) fluctuations, 3D STED could greatly eliminate the out-of-focus signal from the nanotube thus providing more accurate line scan results. Line scan results were analyzed manually only if they showed good signal-to-noise ratio (SNR) and nanotubes were not moving out of focus during the imaging time. All line scan results were analyzed with Matlab systematically to compare with the manually analyzed data.

### Microfluidics

#### Wafer Design and Fabrication

The pattern on the wafer master was designed with AutoCAD 2017. Typically, it consists of an 8 or 12 channel cascade GUV trapping system^36^, with each channel containing 17 GUV traps. The channels are equipped with guiding posts to divert the flow in order to collect GUVs more efficiently into the traps (Fig. S1). For different tasks of the experiment, the microfluidic device is designed with different heights, from 60 to 120 μm. Thus we designed the dimensions of the trapping post in the range of 35-50 μm to meet the desired aspect ratio to prevent the posts from collapsing. A gap size of 10 μm between the posts was used in the system as a compromise between solution exchange rate around the GUVs and vesicle trapping efficiency. In order to obtain high resolution patterns on the wafer, a chrome mask with better resolution was employed. As 5-10 μm is around the resolution limit film masks can offer, we chose chrome masks which can typically provide a resolution down to 1 μm to produce the best pattern and ensure the small post gap on the wafer is retained.

The wafers were produced with conventional soft lithography techniques^36, 50^. A 4-inch diameter silicon wafer was heated and dehydrated at 200 °C on a hotplate for 20 mins to enhance the photoresist adhesion. Then, SU 8-3050 (MicroChem) photoresist was spin coated onto the silicon wafer at a specific rotation speed to obtain the desired thickness. Afterwards, it was pre-baked at 65 °C for 5 mins, 95 °C for 25 mins, and again 65 °C for 5 mins. A mask aligner (EVG-620, EV Group) was employed for UV exposure at 250 mJ/cm^2^ through the mask and therefore writing the desired pattern on the coated wafer. The wafer was then immediately post-baked at 65 °C for 5min, 95 °C for 10 min, and again 65 °C for 5 min. SU-8 developer (Sigma-Aldrich) was used for dissolving the unwanted portion of photoresist in order to obtain the final pattern on the wafer, followed by rinsing the wafer with excess amounts of isopropanol and dried gently with a nitrogen gun. The height of the features was measured by a white-light interferometer (Wyko NT1100). In order to extend its lifespan, the wafer was salinized before use to prevent PDMS adhering to it after curing. It was placed alongside with a few drops of the salinizing agent in a petri dish and put in a vacuum desiccator for 1 min. Then, the wafer was left in the sealed salinizing environment for 24 hours.

#### Chip Fabrication

Microfluidic device was produced by pouring degassed PDMS precursor and curing agent (Sylgard 184, Dow Corning GmbH), at a mass ratio of 10:1, onto the wafer and baking at 80 °C for 2 hours. The PDMS block was gently peeled off from the wafer and cut into pieces with a razor. Inlet and outlet holes were punched with a biopsy punch with a plunger system (Kai Medical). Glass coverslips were cleaned by detergent, rinsed with water and ethanol, blow dried with nitrogen, and baked at 100 °C for 10 min. The PDMS device and coverslip were then plasma-treated for 1 min using high power expanded plasma cleaner (Harrick Plasma). Immediately after plasma treatment, the PDMS device was bonded to the coverslip. The device was further baked at 80 °C for 30 min to accelerate the chemical reaction. The microfluidic devices were stored in a closed box until use. Solution exchange in the microfluidic device was controlled by NeMESYS syringe pump using a 0.5 mL Hamilton gas-tight syringe with appropriate connectors and tubing. Before experiment, the desired amount of solution was filled into the microfluidic device by centrifugation at 900 relative centrifugal force (Rotina 420R, Hettich).

### Interfacial tension measurement

The interfacial tension between the coexisting dextran-rich and PEG-rich phases was measured using a SITE100 spinning drop tensiometer (Krüss GmbH, Hamburg). Circa 1 μL of the PEG-rich droplet was injected into a transparent glass capillary which was prefilled with degassed denser solution of the dextran-rich phase. The horizontally aligned capillary rotated at a speed ω between 500 and 12000 rpm, and the lighter droplet became elongated along the axis of rotation. The interfacial tension, σ, between the two phases was calculated from the Vonnegut equation at a sufficiently high rotation speed when the length of the droplet exceeded 4 times its equatorial diameter. The equation has the form 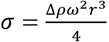, where ∆*ρ* is the density difference between the coexisting phases and *r* is the equatorial radius of the ellipsoidal droplet. The radius was measured in the software by calibrating the pixel size using a stiff cylindrical stick with a known diameter placed in the capillary filled with the same solution of the dextran-rich phase.

### Automated Image Analysis

The peak-detection in tube line scans was accomplished using the findpeaks function of Matlab 2017a (Mathworks Inc.). In principle, the nanotube diameter is calculated from the location of the two high intensity peaks from the nanotube lipid bilayer sidewalls. To identify valid (in focus, revealing the true nanotube diameter) measurements, we considered the additional information of the fluorophore intensity. Because of the asymmetric shape of the STED volume, in which the signal of the fluorophore is integrated, the measured intensity is not constant at different positions along the nanotube cross section. In general, the overlap between the nanotube walls and STED volume is the highest for line scan passing through the maximal tube cross section. Consequently, only measurements with peak intensity above a threshold were considered for analyses. Additionally, the two characteristics peaks of the nanotube walls are required to be within 10% of their respective intensities. Finally, we aimed to filter out small intensity fluctuations which might be misinterpreted as a peak by considering the prominence of the peak to be above a second threshold (see Fig. S6a). To identify the optimal values for the two threshold parameters, we obtained about 500 STED-scans on the nanotubes of a single GUV, where we can safely assume that every nanotube has the same diameter. Now both threshold parameters were varied systematically, to find the parameter-set, which correspond to the lowest standard deviation of the nanotube diameter estimate. The resulting individual measurements of tube diameter (obtained on a single GUV) are shown in Fig. S6b). It is apparent that due to the stochastic nature of the tube movement and measurement errors, not a single tube diameter, but a distribution is obtained (Fig. S6c). Individual data points are discrete corresponding to the finite pixel-size of the measurement (20 nm). Virtually all measurements of this particular nanotube fall in the range of 80 +/-20nm, which is within the resolution limit set by the pixel size and is in correspondence with the estimate of the STED resolution. Note that the actual accuracy as measured by the standard error exceeds the STED resolution (78 nm, S.E. of 4 nm, N=32). The parameters of peak and prominence thresholds obtained in this way were used in the further analysis of nanotube diameters on varying vesicles and varying experimental conditions.

## ASSOCIATED CONTENT

### Supporting Information

Figures with supporting information and movies. This material is available free of charge via the Internet at http://pubs.acs.org.

### Author Contributions

RD, RL and ZZ designed the experiments. RD and ZZ wrote the manuscript with contributions from other coauthors. ZZ performed the experiments. ZZ and DR performed the STED alignment and resolution experiments. JS designed the automated image analysis. TR and ZZ designed the microfluidic chip. All authors have given approval to the final version of the manuscript.

### Notes

The authors declare no competing financial interest.

## ACKNOWLEDGEMENT

This work is part of the MaxSynBio consortium which was jointly funded by the Federal Ministry of Education and Research of Germany and the Max Planck Society.

